# Independence of HIF1a and androgen signaling pathways in prostate cancer

**DOI:** 10.1101/848424

**Authors:** Maxine GB Tran, Becky AS Bibby, Lingjian Yang, Franklin Lo, Anne Warren, Deepa Shukla, Michelle Osborne, James Hadfield, Thomas Carroll, Rory Stark, Helen Scott, Antonio Ramos-Montoya, Charlie Massie, Patrick Maxwell, Catharine ML West, Ian G. Mills, David E. Neal

## Abstract

Androgen signaling drives prostate cancer progression and is a therapeutic target. Hypoxia/HIF1a signaling is associated with resistance to hormone therapy and a poor prognosis in patients treated with surgery or radiotherapy. It is not known whether the pathways operate in cooperation or independently. Using LNCaP cells with and without stable transfection of a HIF1a expression vector, we show that combined AR and HIF1a signaling promotes tumor growth *in vitro* and *in vivo*, and the capacity of HIF1a to promote tumor growth in the absence of endogenous androgen *in vivo*. Gene expression analysis identified 7 genes that were upregulated by both androgen and HIF1a. ChIP-Seq analysis showed that the AR and HIF/hypoxia signaling pathways function independently regulating the transcription of different genes with few shared targets. In clinical datasets elevated expression of 5 of the 7 genes was associated with a poor prognosis. Our findings suggest that simultaneous therapeutic inhibition of AR and HIF1a signaling pathways should be explored as a potential therapeutic strategy.

## 1. Introduction

Androgen signaling drives prostate cancer development and progression. Endogenous androgens, testosterone and dihydrotestosterone, bind to the intracellular androgen receptor (AR) which translocates to the nucleus. The AR functions as a transcription factor and activates downstream signaling pathways associated with proliferation, invasion and metabolism [1]. Androgen deprivation therapy (ADT) inhibits AR signaling by blocking the production of androgens or by inhibiting androgen binding to the AR. ADT is used to treat localized, locally advanced and metastatic disease and an estimated 50% of prostate cancer patients receive ADT [2]. Although ADT is initially effective, resistance subsequently develops and the AR signaling pathway remains active even in the absence of endogenous androgens. The development of androgen independent or castrate-resistant prostate cancer (CRPC) is associated with the presence of metastases and a rapid clinical demise [3]. Identifying which patients will progress to CRPC is a major challenge in the treatment of prostate cancer. Understanding the biology that underpins progression to CRPC will support the development of novel strategies to identify, prevent and treat CRPC.

Mechanisms thought to contribute to AR activity in CRPC include gene amplification, activating mutations and cross-talk with other signaling pathways such as the hypoxia inducible factor (HIF) pathway [4]. HIF is a heterodimer, consisting of a constitutively stable HIF1b and a tightly regulated HIF1a subunit. Under oxygenated conditions, the HIF1a protein is ubiquitinated and rapidly degraded. In the absence of oxygen, HIF1a is stabilized and dimerizes with HIF1b subunits to form an active HIF transcription complex. HIF translocates to the nucleus and induces the expression of genes associated with metabolism, angiogenesis, invasion and cell survival. Hypoxia-independent stabilization can also occur– a condition referred to as pseudohypoxia [5] [6]. Expression of HIF is associated with increased risk and a poor prognosis in prostate cancer [7, 8].

Crosstalk between the AR and hypoxia/HIF has been reported. ADT in hypoxia promotes adaptive androgen/AR-independence, and confers resistance to androgen/AR-targeted therapy (Geng et al 2018). Co-immunoprecipitation assays have confirmed a direct interaction between AR and HIF1a, and ChIP analysis showed HIF1a interacts with the AR on the PSA gene promoter [9]. Hypoxia induced activation of HIF can also increase expression of the AR [10, 11]. As AR and HIF signaling pathways are major signaling hubs and oncogenic drivers of prostate cancer progression, this study aimed to investigate further the relationship between them. Here we report for the first time that combined AR and HIF1a signaling *in vivo* promotes tumor growth and demonstrate the capacity of HIF1a to promote tumor growth in the absence of endogenous androgen *in vivo*. We also show that the AR and HIF/hypoxia signaling pathways function independently regulating the transcription of different subsets of genes with few shared targets.

## 2. Materials and methods

### 2.1 Cell culture

LNCaP, LNCaP-Bic, LNCap-OHF and PC3 cell lines (and the corresponding stable transfectants) were cultured in RPMI with glutamine and 10% fetal calf serum. For hypoxia experiments, cells were exposed to 1% oxygen using either a hypoxic workstation (INVIVO2, Ruskinn, Leeds, UK) or a hypoxic incubator. For AR signaling experiments, LNCaP cells were grown in charcoal stripped serum for 96 h prior to adding synthetic androgen (R1881) or vehicle control (ethanol).

### 2.2 Infection of HIF1a retroviral vectors

A model of pseudohypoxia was established in androgen-sensitive LNCaP cells by viral transfection of a vector encoding HIF1a with two amino acid substitutions which prevented its degradation in the presence of oxygen. Viral supernatants were prepared by transfecting the Phoenix packaging cell line (Orbigen, San Diego, CA) using Lipofectamine 2000 (Life Technologies, Paisley, UK). After initial transfection, Phoenix cells were grown at 32°C. The supernatant was collected and filtered (0.45 μm), then supplemented with a 1:4 volume of fresh medium with 7.5 μg/mL Polybrene (Sigma, Poole, UK), and added to LNCaP cells plated on p100 dishes at 30-40% confluence. After 20 h, cells were washed, and fresh media added for 20 h before a second round of transfection and G418 selection. The constitutively active form of HIF1a (carrying two substitutions: P402A and P564A) was cloned into pBMN-I-EGFP.

### 2.3 Western blot analysis

Cell lysis involved urea-SDS buffer supplemented with phenylmethylsulfonyl fluoride (PMSF) as previously described [1]. Immunoblots were visualized with enhanced chemiluminescence reagent or enhanced chemiluminescence plus reagent (Amersham, Arlington Heights, IL). Antibodies used were HIF1a (clone 54, Transduction Labs, Lexington KY) and α tubulin (CRUK).

### 2.4 Xenograft experiments

Xenograft tumors were generated with LNCaP/Empty and LNCaP/HIF1-clone 1 cells that stably expressed a fusion protein of luciferase and yellow fluorescent protein (YFP). There were four groups (Full/non-castrated + LNCaP/Empty, Full/non-castrated + LNCaP/HIF1a clone 1, Castrated + LNCaP/Empty and Castrated + LNCaP/HIF1a clone 1), each consisting of five mice. Tumor growth was monitored weekly through bioluminescence with an IVIS camera (Xenogen) [1].

### 2.5 Clinical material and immunohistochemistry

Clinical samples were collected from Cambridge University Hospitals NHS Trust as part of the PROMPT study, and ethical approval was granted by the local research and ethics committee (LREC number: 02/281M) and by the multicenter research and ethics committee (MREC number 01/4061). A tissue microarray (TMA) was constructed, consisting of at least two tumor cores with matched benign prostate tissue cores from each of 41 patients with CRPC. Sections 3 μm thick were mounted on Snowcoat X-tra slides (Surgipath, Richmond, IL), dewaxed in xylene and rehydrated using graded ethanol washes. For antigen retrieval, sections were immersed in preheated DAKO target retrieval solution and treated for 90 s in a pressure cooker (DAKO, Glostrup, Denmark). Antigen/antibody complexes were detected using the DAKO catalyzed signal amplification (CSA) system according to the manufacturer’s instructions. Sections were counterstained with hematoxylin for 30 s, dehydrated in graded ethanol washes and mounted (Lamb, London, UK). A rabbit pAb was used for HIF1a immunohistochemistry in the xenograft tumors (#NB 100-479, Novus Biologicals, Oxford, UK) and a mouse mAb was used for HIF1a immunohistochemistry in the CR-TMA (H1α67 # NB 100-105, Novus Biologicals). Immunohistochemistry staining was scored by two independent blinded assessors as 1 (negative), 2 (<25% of nuclei staining), 3 (25-50% of nuclei staining), 4 (majority of cells – weak staining), 5 (majority of cells – moderate staining) and 6 (majority of cells – strong staining).

### 2.6 Illumina HumanWG v2 BeadArray data analysis

LNCaP, LNCaP/Empty and LNCap/HIF1a clone 1 cells were grown in charcoal stripped serum for 96 h prior to adding 1 nM R1881 or 0.01% ethanol (vehicle control) for 4 h and extracting RNA. For hypoxia experiments, LNCaP cells were exposed to 1% hypoxia or normoxia for 24 h prior to RNA extraction. Gene expression data were generated using the Illumina HumanWGv2 BeadArrays. After background correction, normalization and log2 transformation, differential expression analysis was performed with LIMMA on probe set level. Probe sets with bad and no match probe scores were omitted from analysis. False discovery rate adjusted *P* value of 0.05 and 1.5 fold change were applied as cut-off.

### 2.7 Chromatin Immunoprecipitation

ChIP and re-ChIP was performed as previously described [12, 13]. Cells were cultured in phenol red-free RPMI media supplemented with 10% charcoal dextran-stripped FBS for 72 h before adding 1 nM R1881 or 0.01% ethanol for 4 h. For hypoxic experiments, cells were placed in a hypoxic incubator at 1% oxygen for 12 h prior to adding R1881 or ethanol. AR (AR N20, Sc-816X, Santa Cruz), HIF1a (ab2185, Abcam), H3K4me1 (pAb 194-050 Diagenode) and H3K4me3 (pAb 003-050, Diagenode, Seraing, Belgium) antibodies were used in the assay. ChIP enrichment was tested by real-time PCR and the remainder was used for single-end SOLEXA library preparation.

### 2.8 ChIP-seq SOLEXA library preparation

Single-end SOLEXA sequencing libraries were prepared as previously described [13]. Sequence reads were generated using an Illumina Genome Analyzer II and mapped to the reference human genome before peak calling. Called peaks were analysed in R using ChIPpeakAnno package [14].

### 2.9 Data deposition

Microarray and ChIP-seq data generated have been deposited within the National Center for Biotechnology Information (NCBI) Gene Expression Omnibus (GEO) database (https://www.ncbi.nlm.nih.gov/geo/) under GSE114734.

### 2.10 Patient cohorts, endpoints and statistical analysis

Five prostate cancer gene expression cohorts with publically available patient survival data were used to evaluate the prognostic significance of selected genes: TCGA [15], Taylor et al GSE21032 [16], Long et al GSE54460 [17], Ross-Adams et al GSE70770 [15] and Sboner et al GSE16560 [18] (Supplementary Table I). For TCGA, GSE54460, GSE70770, GSE16560 gene expression data were downloaded directly. For GSE21032, raw CELL files were downloaded and processed using aroma package.

Biochemical recurrence free (BCR) survival was the primary endpoint, except for Sboner where only overall survival was available. Patients were stratified into high and low groups based on cohort median expression of the gene of interest. Survival estimates were performed using the Kaplan-Meier method. The log-rank test was used to test the null hypothesis of equality of survival distributions. Hazard ratios (HR) and 95% confidence intervals (CI) were obtained using the Cox proportional hazard model.

## 3. Results

### 3.1 HIF1a expression promotes proliferation and resistance to ADT in vitro and in vivo

The stable overexpression of HIF1a in LNCaP/HIF1a clone 1 and clone 2 (Supplementary Fig 1A) cells increased proliferation and resistance to ADT (bicalutamide) *in vitro* (Fig 1). Growth rate decreased in response to ADT in LNCaP/Empty but not LNCaP/HIF1a cells. HIF1a expression was also detected in normoxia in ADT resistant (LNCaP-Bic, LNCaP-OHF) and androgen-independent (PC3) cells but not in androgen sensitive (LNCaP) cells (Supplementary Fig 1). LNCaP/HIF1a xenografts grew faster than the LNCaP/Empty tumors, and were resistant to ADT (castrated mouse model; Fig 2).

**Figure 1.**
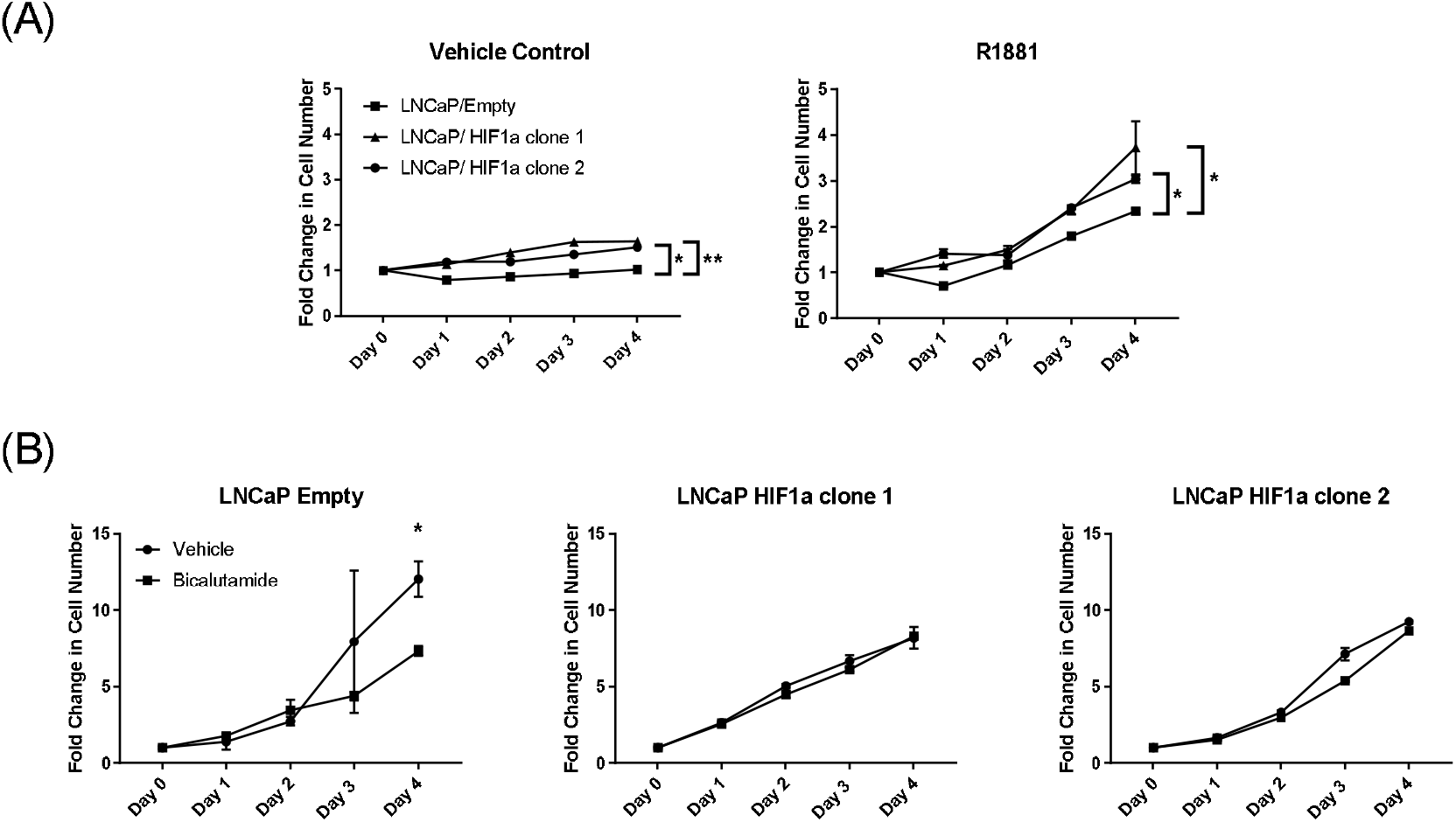
HIF1a overexpression in the androgen dependent LNCaP cell line increased proliferation and resistance to androgen deprivation therapy. A, stable HIF1a expression increased cell proliferation compared to the LNCaP/Empty control cells when cells were treated with the ethanol vehicle control or synthetic androgen R1881 (two way multiple comparison ANOVA; *p<0.05, **p<0.01). B, stable HIF1a expression led to resistance to bicalutamide treatment (two-tail t-test, * p=0.058). Data points represent the mean of three intra-assay and two biological repeats ± SEM. Differences in growth rates are due to cells grown in media containing charcoal stripped serum (androgen depleted media) for 96h prior to treatment (A) or in standard RPMI median with FBS that contains androgen and growth stimulants (B).

**Figure 2.**
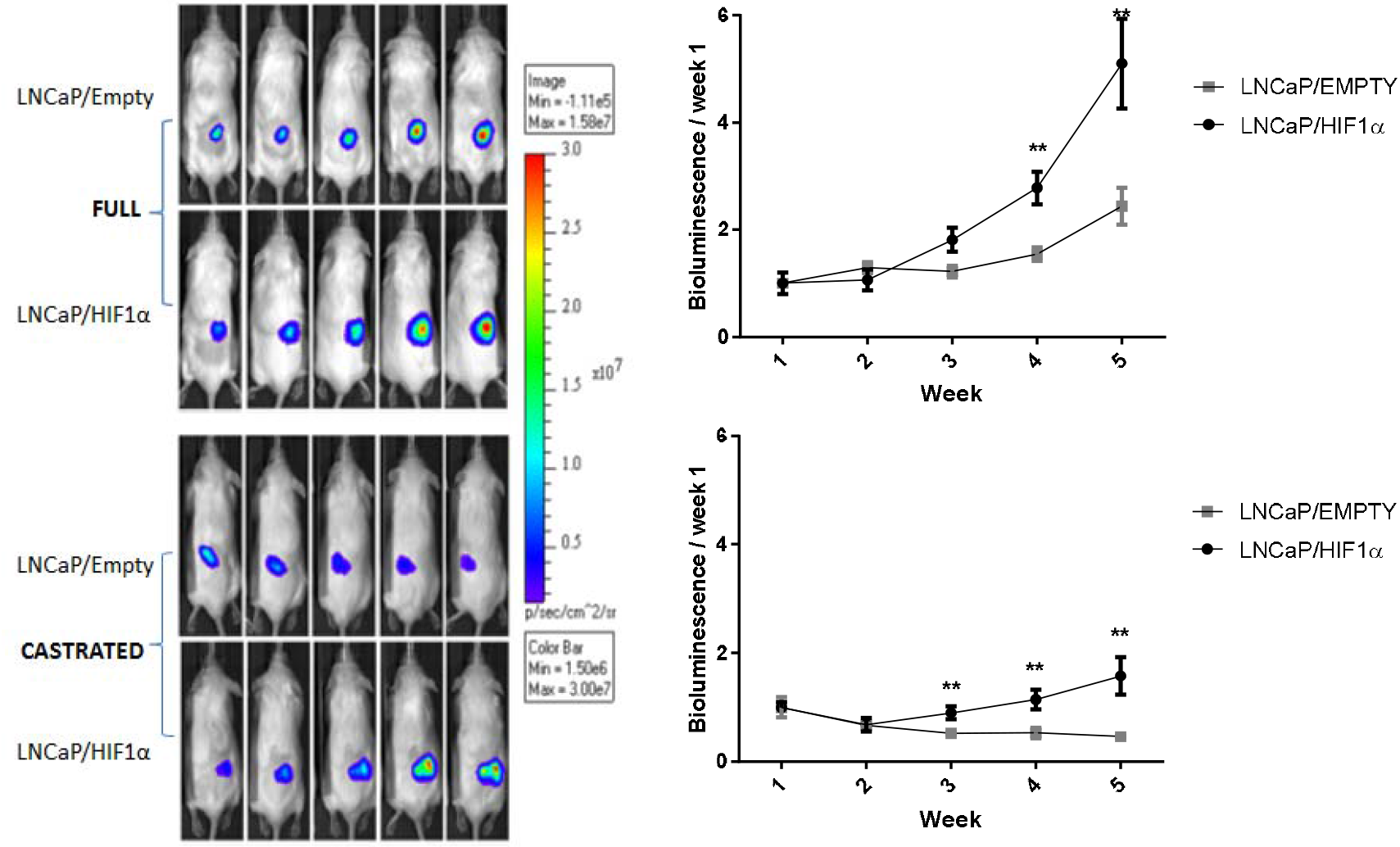
HIF1a accelerated tumor growth in non-castrated (full) and castrated mice. A, tumor xenografts derived from the LNCaP/HIF1a clone 1 cell line showed accelerated growth in full and castrated mice however LNCaP/HIF1a tumors grew significantly slower in the castrated mouse compared to the full mouse model. In the castrated mice LNCaP/HIF1a tumors continued to grow whilst the empty control clone regressed (data points represent the mean ± SEM, two-way ANOVA; **p<0.01).

### 3.2 Castrate resistant prostate cancers have high HIF1a expression

HIF1a immunohistochemistry showed high levels of HIF1a expression in CRPC. All CRPC biopsies expressed HIF1a in comparison with only 57% of benign tumors. Sixty-eight percent of CRCP biopsies had strong HIF1a staining compared with just 8% of benign tissue (Supplementary Fig 2).

### 3.3 Only seven genes upregulated by both androgen and HIF1a

In LNCaP cells, gene expression analysis identified 336 genes upregulated in response to synthetic androgen (Supplementary Table II) and 579 genes in response to hypoxia (Supplementary Table III). Forty-seven genes were upregulated by both androgen and hypoxia (Fig 3A). There were 149 genes upregulated in response to synthetic androgen in LNCaP/Empty cells and 56 genes upregulated in response to stable HIF1a expression (Fig 3B). Only seven genes (*TWIST1, KCNN2, PPFIBP2, JAG1, SPRED1, IGFBP3, NDRG1*) were upregulated by both androgen and HIF1a (Fig 3B). Three genes (*SPRED1, IGFBP3, NDRG1*) were upregulated by androgen, hypoxia and stable HIF1a expression.

**Figure 3.**
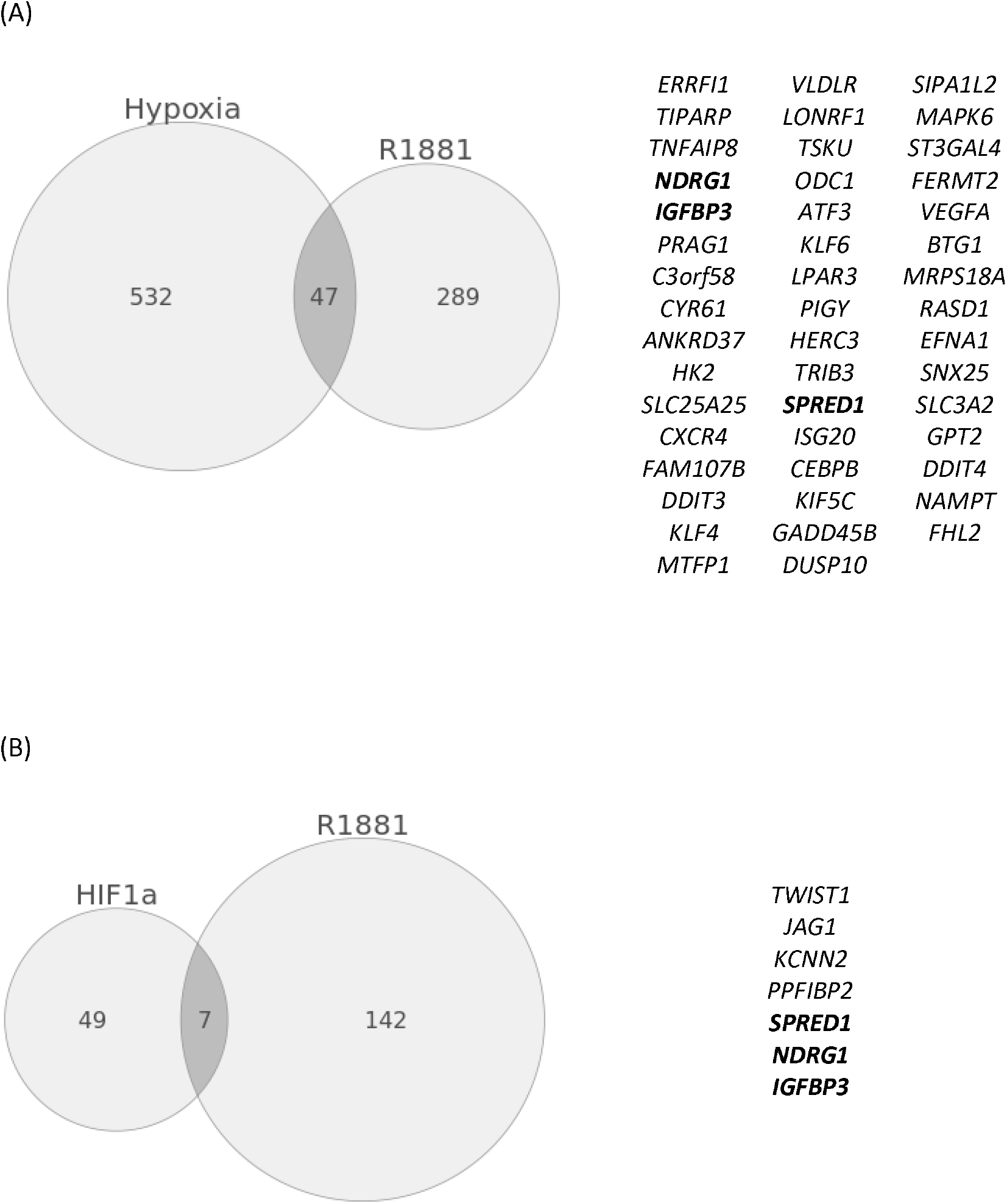
Genes upregulated by androgen (R1881), hypoxia and HIF1a in LNCaP cells. A, 47 genes upregulated by androgen (LNCaP vehicle control vs. LNCaP R1881, right circle) were independently upregulated by hypoxia (LNCaP normoxia vs. LNCaP 1% hypoxia, left circle). B, 7 genes upregulated by HIF1a overexpression (LNCaP Empty vs. LNCaP HIF1a, left circle) were also independently upregulated by androgen (LNCaP Empty vehicle control vs. LNCaP Empty R1881, right circle). Three genes were independently upregulated by and androgen, hypoxia and HIF1a *(SPRED1, NDRG1* and *IGFBP*).

### 3.4 AR and HIF transcription factor binding sites increase differently in response to androgen and hypoxia, respectively

ChIP-seq analysis identified global AR and HIF DNA binding sites in LNCaP cells exposed to synthetic androgen under normoxia and hypoxia. There were more AR (called peaks range 18,404 to 70,064) compared to HIF (range 523 to 5,795) transcription factor binding sites (Table I). The number of AR binding sites increased with androgen treatment and decreased in hypoxia. However, while hypoxia almost halved the number of AR binding sites, they increased when androgen was added under hypoxia (from 18,404 to 45,635) suggesting an interplay between hypoxia and androgen signalling at the level of AR recruitment to chromatin. AR binding sites were highly conserved between the vehicle control and androgen treated cells exposed to normoxia (86%) and hypoxia (79%) (Fig 4A). As expected hypoxia increased the number of HIF binding sites, the greatest number of HIF binding sites was observed in cell treated with combined hypoxia and androgen treatment (Table I). HIF binding sites were not conserved in cells between normoxia and hypoxia in the absence (6%) or presence (3%) of androgen (Fig 4B). These experiments show androgen increases the number of AR binding sites while conserving those present in the absence of androgen, i.e. AR binds to additional sites. In contrast, hypoxia increases the number of HIF binding sites but with little conservation, i.e. HIF binds to different sites in hypoxia. These observations highlight differences in the effect on transcription when the AR responds to androgen and HIF to hypoxia.

**Figure 4.**
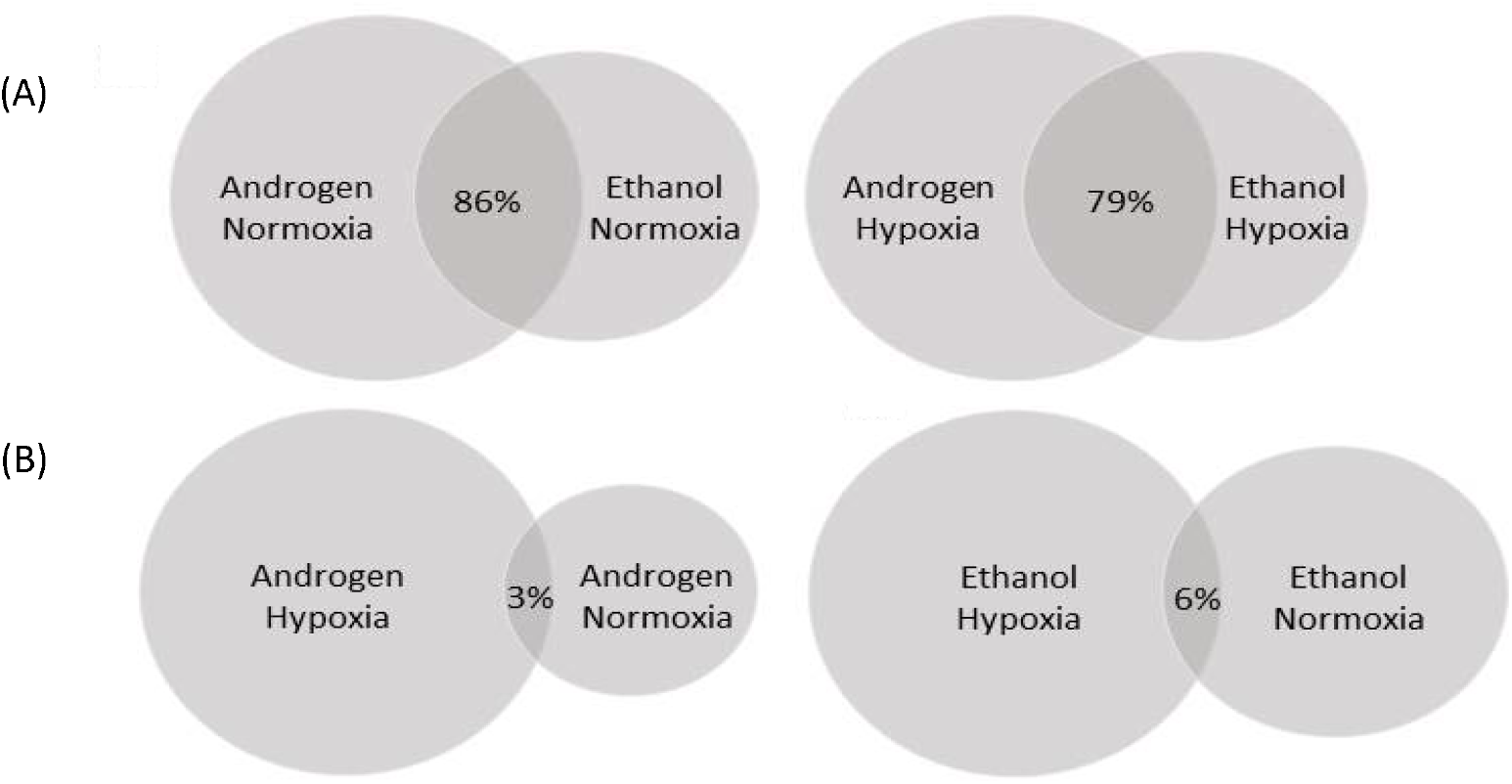
Conservation of AR and HIF binding sites. A, The majority of AR binding sites were conserved following androgen treatment. Of the 35320 AR called peaks in the normoxic ethanol vehicle control 86% were conserved in the normoxic R1881 treated cells (*left*). Of the 18404 AR called peaks in the hypoxic ethanol vehicle control 79% were conserved in the hypoxic androgen treated cells under (*right*). B, HIF binding sites were not conserved upon hypoxic exposure. Of the 523 HIF called peaks in the normoxic androgen cells 3% were conserved in the hypoxic androgen treated cells under (*left*). Of the 1181 HIF called peaks in the normoxic ethanol vehicle control 6% were conserved in the hypoxic ethanol vehicle control cells (*right*).

**Table I.** ChIP-seq global called peaks for AR, HIF and histone methylation markers.

| Antibody | Condition | Called peaks |
| --- | --- | --- |
| AR | R1881/Normoxia | 70064 |
| AR | R1881/Hypoxia | 45635 |
| AR | Ethanol/Normoxia | 35320 |
| AR | Ethanol/Hypoxia | 18404 |
| HIF | R1881/Normoxia | 523 |
| HIF | R1881/Hypoxia | 5795 |
| HIF | Ethanol/Normoxia | 1181 |
| HIF | Ethanol/Hypoxia | 1746 |
| H3K4Me1 | R1881/Normoxia | 74269 |
| H3K4Me1 | R1881/Hypoxia | 79415 |
| H3K4Me1 | Ethanol/Normoxia | 90818 |
| H3K4Me1 | Ethanol/Hypoxia | 95424 |
| H3K4Me3 | R1881/Normoxia | 16419 |
| H3K4Me3 | R1881/Hypoxia | 18627 |
| H3K4Me3 | Ethanol/Normoxia | 24358 |
| H3K4Me3 | Ethanol/Hypoxia | 38929 |

Not all binding sites reflect sites of transcriptional activity. A comparison with histone marks can refine this landscape and help to identify those sites most likely to be active: H3K4me1 is enriched at active and primed enhancers and H3K4me3 in a promoter (i.e. most likely to be active) and stable H3K4me3 has been associated with transcription initiation [19, 20]. Table I shows androgen globally decreased both histone markers. In comparison, hypoxia globally increased the number of histone marker binding sites. Fig 5 A-D shows no change in the distribution of H3K4me1 in response or androgen or hypoxia. Fig 5 E-H shows an increase in the percentage of H3K4me3 located within promoter regions in response to androgen and a decrease in response to hypoxia. These observations are consistent with androgen increasing and hypoxia decreasing transcription initiation at transcriptional start sites.

**Figure 5.**
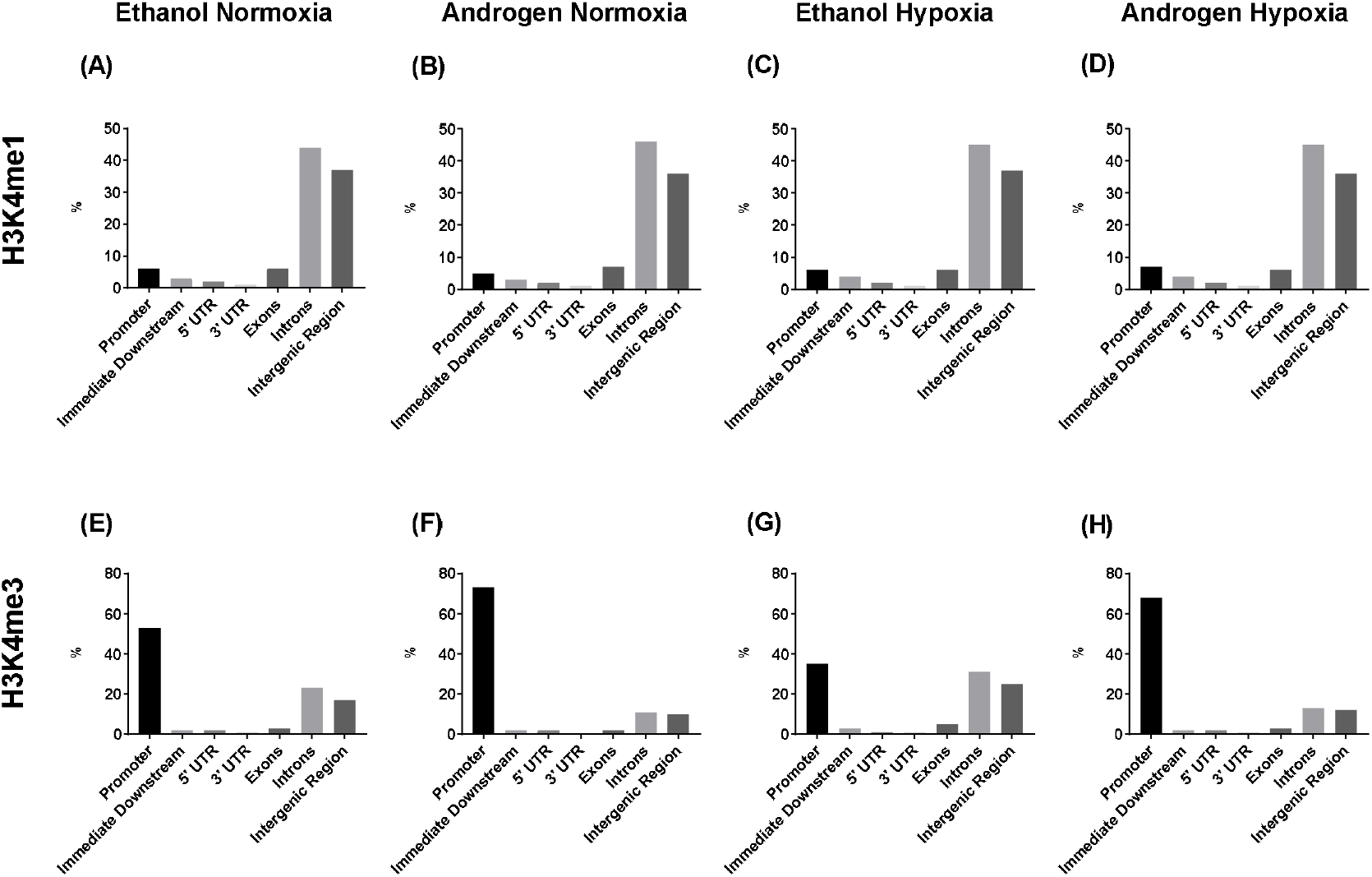
The global genomic distribution of the histone markers, H3K4me1 and H3K4me3, in the LNCaP ChIP-seq analysis. The distribution of the H3K4me1 marker did not change with androgen treatment or hypoxia (A-D). Hypoxia decreased H3K4me3 markers within promoter regions (E vs G). Synthetic androgen R1881 increased the location of H3K4me3 markers within promoter regions under normoxia (E vs F) and hypoxia (G vs H).

The locations of the transcription factors (AR, HIF) and the histone markers (H3K4me1, and H3K4me3) were analysed within the exons and introns of the seven genes identified in the gene expression analysis (Supplementary Fig 3). Neither AR nor HIF bound within the *TWIST1* and *IGFBP3* genes (data not shown). There were more AR, HIF, H3K4me1 and H3K4me3 binding sites in *KCNN2* and *PPFIBP2* compared to the other genes (Table II). These observations suggest that KCNN2 and PPFIBP2 are directly regulated by promoter proximal and intragenic recruitment of the AR and HIF1 whereas TWIST1 and IGFBP3 may be enhancer regulated. Indeed changes in IGFBP3 expression have been shown to be affected by and to affect the expression of a range of genes through long-range chromatin and interchromosomal interactions [21]. In addition, TWIST1 is known to function as a transcriptional driver of EMT. Consequently, although the number of genes we have identified as co-ordinately regulated by the AR and HIF1 is small in number their impact may be far-reaching.

**Table II.** Numbers of binding sites of transcription factors and histone markers in selected gene in LNCap cells

|  | *KCNN2* | *PPFIBP2* | *JAG1* | *SPRED1* | *NDRG1* |
| --- | --- | --- | --- | --- | --- |
| AN-H-AR | 4 | 9 | 1 | 1 | 0 |
| AN-H-HIF | 0 | 4 | 0 | 0 | 2 |
| AN-H-ME1 | 13 | 25 | 2 | 2 | 6 |
| AN-H-ME3 | 0 | 2 | 1 | 1 | 2 |
| AN-N-AR | 8 | 11 | 2 | 6 | 3 |
| AN-N-HIF | 1 | 0 | 0 | 0 | 0 |
| AN-N-ME1 | 18 | 15 | 1 | 6 | 4 |
| AN-N-ME3 | 1 | 3 | 1 | 2 | 1 |
| ET-H-AR | 1 | 7 | 1 | 0 | 0 |
| ET-H-HIF | 0 | 0 | 0 | 0 | 0 |
| ET-H-ME1 | 15 | 19 | 2 | 0 | 5 |
| ET-H-ME3 | 4 | 8 | 2 | 1 | 4 |
| ET-N-AR | 6 | 9 | 1 | 3 | 1 |
| ET-N-HIF | 0 | 0 | 0 | 0 | 0 |
| ET-N-ME1 | 18 | 11 | 2 | 2 | 5 |
| ET-N-ME3 | 4 | 5 | 1 | 1 | 1 |

AN, synthetic androgen (R1881) treatment; ET, ethanol vehicle control; N, normoxia; H, hypoxia; AR=androgen receptor; ME1, H3K4me1; ME3, H3K4me3.

### 3.5 Effect of TWIST1, KCNN2, PPFIBP2, JAG1, SPRED1, IGFBP3 and NDRG1 on prognosis

Five publically available prostatectomy gene expression cohorts were used to test the prognostic significance of the seven genes upregulated by androgen, stable HIF1a expression and hypoxia (Table III). *TWIST1* was the most prognostic with high expression associated with poor a prognosis in three cohorts. Five of the genes were prognostic in a single cohort and *SPRED1* had no prognostic significance (Table III). We further compared *TWIST1* to a recently published hypoxia-gene associated prognostic signature for prostate cancer [22]. The 28-gene prognostic signature was derived from the TCGA cohort, and had a significant proportion of genes absent in Sboner et al cohort. In Taylor et al both *TWIST1* (HR 2.45, 95% CI 1.01-5.93, P=0.047) and the 28-gene signature (HR 4.48, 95% CI 1.67-12.04, P=0.0030) retained prognostic significance.

**Table III.** Prognostic significance of selected genes in prostate cancer cohorts.

|  | Swedish | Cambridge | Long | Taylor | TCGA |
| --- | --- | --- | --- | --- | --- |
| *TWIST1* | **2.05**<br>**(1.30-3.22)**<br>**P=0.0019** | 0.95<br>(0.39-2.35)<br>P=0.917 | 1.46<br>(0.84-2.54)<br>P=0.178 | **2.69**<br>**(1.11-6.49)**<br>**P=0.028** | **1.72**<br>**(1.13-2.60)**<br>**P=0.01** |
| *KCNN2* | **2.39**<br>**(1.51-3.79)**<br>**P=0.000216** | 1.91<br>(0.75-4.86)<br>P=0.17 | 1.52<br>(0.87-2.27)<br>P=0.14 | 1.78<br>(0.78-4.08)<br>P=0.17 | 0.84<br>(0.56-1.26)<br>P=0.41 |
| *SPRED1* | NA | 0.90<br>(0.36-2.23)<br>P=0.81 | 0.76<br>(0.44-1.32)<br>P=0.33 | 0.96<br>(0.43-2.13)<br>P=0.91 | 0.96<br>(0.64-1.44)<br>P=0.85 |
| *PPFIBP2* | NA | 1.21<br>(0.49-3.02)<br>P=0.68 | **1.73**<br>**(0.99-3.03)**<br>**P=0.05** | 0.86<br>(0.38-1.92)<br>P=0.70 | 0.76<br>(0.50-1.13)<br>P=0.17 |
| *JAG1* | 1.36<br>(0.88-2.10)<br>P=0.17 | 2.23<br>(0.85-5.88)<br>P=0.10 | **2.293**<br>**(1.28-4.12)**<br>**P=0.005** | 1.25<br>(0.56- 2.80)<br>P=0.59 | 1.30<br>(0.87-1.94)<br>P=0.21 |
| *IGFBP3* | 1.00<br>(0.65-1.55)<br>P=0.99 | 1.09<br>(0.44-2.68)<br>P=0.86 | 1.56<br>(0.89-2.73)<br>P=0.12 | 1.99<br>(0.85-4.66)<br>P=0.11 | **2.07**<br>**(1.36-3.20)**<br>**P=0.0008** |
| *NDRG1* | **2.28**<br>**(1.43-3.64)**<br>**P=0.0005** | 1.43<br>(0.58-3.56)<br>P=0.44 | 0.81<br>(0.47- 1.41)<br>P=0.45 | 0.78<br>(0.34- 1.75)<br>P=0.54 | 0.96<br>(0.64-1.43)<br>P=0.82 |

BCR, biochemical recurrence; OS, overall survival; NA, not applicable.
Values are hazard ratios (95% confidence intervals). Cohorts were stratified by the median expression of each gene.

## 4. Discussion

Hypoxia and HIF1a signaling are widely regarded as cause and consequence, but there is increasing evidence of pseudohypoxia - the expression of HIF1a in normoxia – in multiple cancers [23]. Our LNCaP/HIF1a clones represent a model of pseudohypoxia. Stable HIF1a increased cell growth in the absence and presence of the synthetic androgen R1881, and promoted resistance to ADT *in vitro* and *in vivo*. Hypoxia and HIF have already been implicated in the development and progression of CRPC [24, 25]. Hypoxia was shown to induce AR independence and confer resistance to ADT through a metabolic switch favoring glycolysis [26]. Pseudohypoxia has also been linked to the metabolic switch from oxidative phosphorylation to glycolysis [27]. Expression of HIF1a in normoxia has been reported in androgen dependent prostate cells and in this study we report expression of HIF1a in cells resistant to ADT (LNCaP-Bic, LNCaP-OHF) and in the androgen independent PC3 cell line ^10 22^. This study adds to the evidence implicating hypoxia and HIF1a in androgen independence, CRCP and ADT resistance.

The high expression of HIF1a in CRPC further supports the role of HIF1a in aggressive, androgen dependent prostate cancer. Whether the high expression of HIF1a was associated with pseudohypoxia or hypoxia could not be determined in this study. In future studies the hypoxia marker pimonidazole alongside HIF1a would provide a valuable insight into the contribution of hypoxia and pseudohypoxia in CRPC.

Gene expression analysis showed few genes were regulated in common by AR, hypoxia, and HIF1a. The finding suggests the signaling pathways act independently and regulate the expression of different subsets of genes. Other studies have reported both positive and negative crosstalk between androgen/AR and hypoxia/HIF1a [26, 28, 29]. Globally there were substantially more AR binding sites than HIF binding sites, demonstrating androgen signaling dominance over HIF signaling in the prostate cancer cells studied. Interestingly, hypoxia decreased the number of AR binding sites. This observation contrasts with studies showing hypoxia enhances AR activity [29–31]. The variability in concentration and duration of R1881 treatment and hypoxia across studies is most likely responsible for the conflicting results. The observed decrease in androgen binding sites under hypoxia in our study may be explained by conformational changes in chromatin structure induced by 12h exposure to 1% hypoxia, which may restrict the accessibility of AR binding sites [32].

The locations of the AR DNA binding sites in the ethanol vehicle control and R1881 treated cells were highly conserved. The 2-fold increase in AR binding sites with R1881 treatment added to the existing AR binding sites that were occupied in the absence of R1881. In comparison, the DNA binding sites occupied by HIF under normoxia were located in different regions to the HIF binding sites occupied under hypoxia, indicating that HIF binds to different sites in the DNA and promotes the expression of a different subset of genes under pseudohypoxia and hypoxia. Histone markers associated with active transcription were globally decreased within the DNA following synthetic androgen R1881 treatment. In contrast hypoxia marginally increased the presence the two histone markers, it has previously been reported that hypoxia rapidly increases histone methylation independently of HIF [33]. Despite decreasing the prevalence of H3K4me3, the location of the histone marker within promoter regions was increased as a result of R1881 treatment and indicates enhanced transcriptional activity.

We found few HIF transcription factor binding sites within the introns and exons of the seven genes upregulated by androgen and HIF1a suggesting the HIF regulated expression of these genes is most likely driven by it binding to distal sites [34]. The greatest number of AR binding sites within the genes was observed with androgen treatment under normoxia, with a reduction in the number of AR binding sites under hypoxia. This decrease in AR binding sites under hypoxia was observed globally, possibly as a result of hypoxia induced conformational changes in the DNA which restrict the accessibility of AR binding sites.

Of the seven genes upregulated by both androgen and HIF *TWIST1* was the most prognostic. Upregulated *TWIST1* and AR expression have previously been reported in a castration resistant LNCaP mouse model, implicating crosstalk between epithelial mesenchymal transition and castration resistance [35]. *TWIST1* was also shown to upregulate AR expression and to be upregulated in response to ADT [36]. The variability in prognostic significance between the cohorts may in part be due to use of different gene expression platforms. A further limitation is that most patients in the cohorts had primary prostate cancer treated by radical prostatectomy without hormone therapy and were mostly low and intermediate risk patients. Considering the seven genes identified in this study are upregulated by androgen, HIF1a and/or hypoxia it is hypothesized that they promote disease progression and development of CRPC and it would be interesting to look at the expression of these genes in high risk and advanced prostate cancer cohorts. As AR and HIF signaling axes are active in CRPC these seven genes are potential biomarkers of aggressive disease that might be useful to predict likely disease progression towards CRPC [37, 38].

In this study the absence of HIF1a and endogenous androgen *in vivo* resulted in regression of tumor growth but HIF1a signaling could restore tumor growth in the absence of AR signaling. The data presented here indicate simultaneous therapeutic inhibition of the HIF1a and AR signaling pathways is a potential therapeutic strategy, as has previously been proposed [38]. We show that the oncogenic signaling pathways target the expression of different subsets of genes but both promote proliferation, tumor growth and disease progression. The relationship between the AR and HIF1a signaling pathways and their association with the development of CRPC could be exploited to identify predictive biomarkers of progression to CRPC and dual targeting of the AR and hypoxia/HIF1a should be further investigated for patients most at risk of developing CRPC.

## Acknowledgements

We acknowledge members of the Uro-Oncology laboratory for helpful discussions. We thank research staff of Cancer Research UK Cambridge Institute core facilities for technical support.

