## Supplementary Data for "Independence of HIF1a and androgen signaling pathways in prostate cancer"

(A)

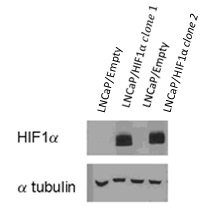

(B)

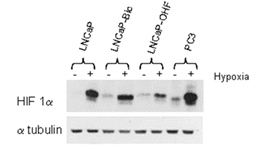

**Supplementary Figure 1.** A, stable overexpression of HIF1a was established in two clones (LNCaP/HIF1a). HIF1a expression was undetectable in the LNCaP/Empty control cell line. B, HIF1a expression was detectable under normxic conditions in the LNCaP bicalutamide resistant (LNCaP-Bic), LNCaP hydroxy flutamide resistant (LNCaP-OHF) and androgen independent PC3 but not in the androgen dependent LNCaP. Exposure to 1% oxygen increased HIF1a expression in all of the cell lines.

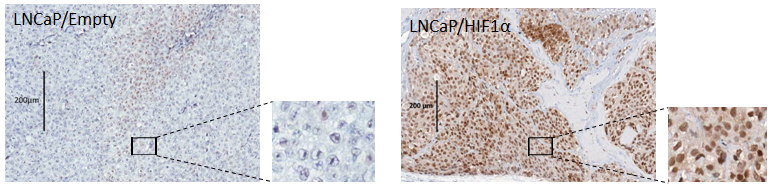

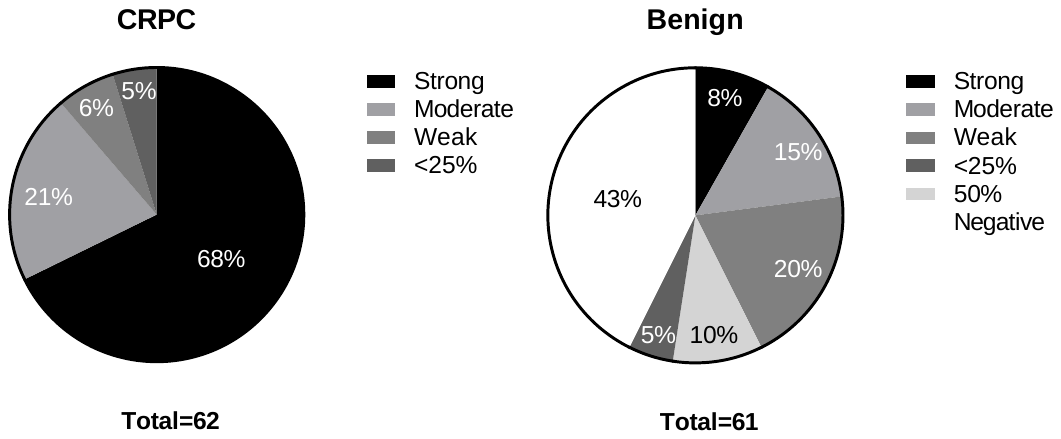

(B)

(A)

**Supplementary Figure 2.** HIF1a immunohistochemistry was performed on the CRPC and matched benign tissue TMAs. A, *ex vivo* immunohistochemistry confirmed HIF1a overexpression in the LNCaP/HIF1a tumors and negative HIF1a staining in the LNCaP/Empty tumors. B, TMAs were constructed using tissue biopsies from 41 patients, 62 tumor and 61 matched benign tissues were stained for HIF1a. Constitutively high expression of HIF1a was observed in CRPC (68%) compared to benign tissue (8%). All of the CRPC tissue stained positive for HIF1a, of the benign tissues 26% were HIF1a staining negative.

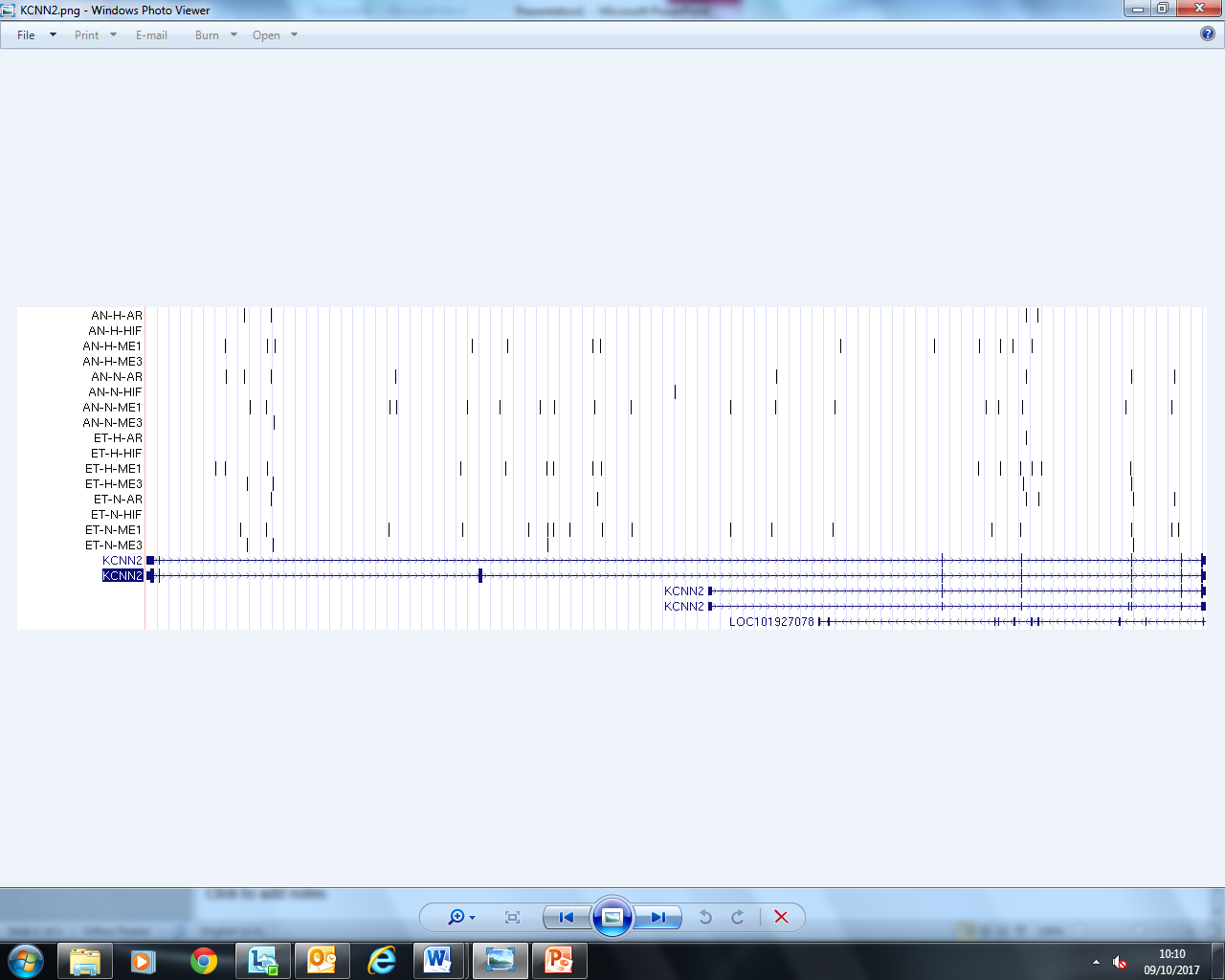

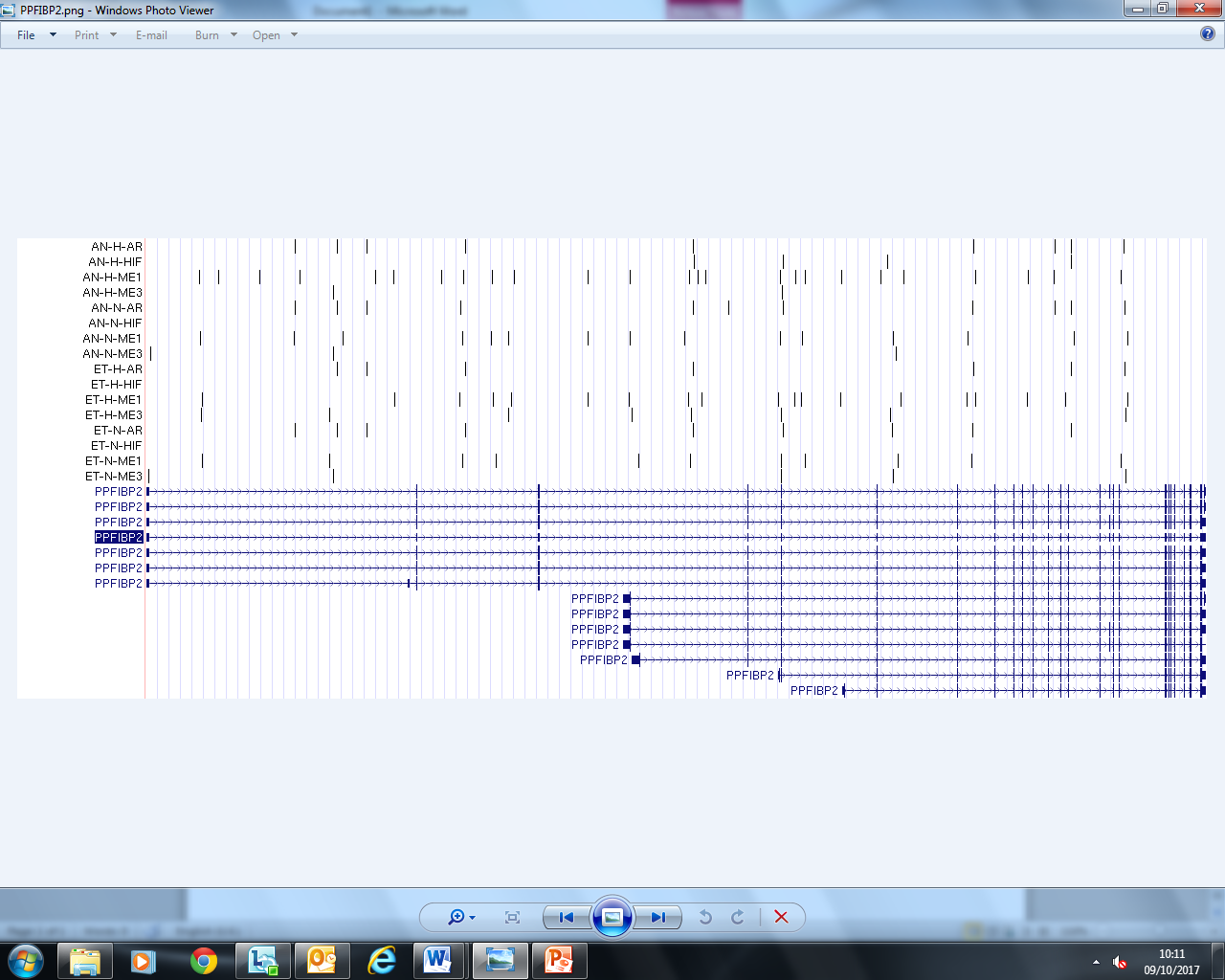

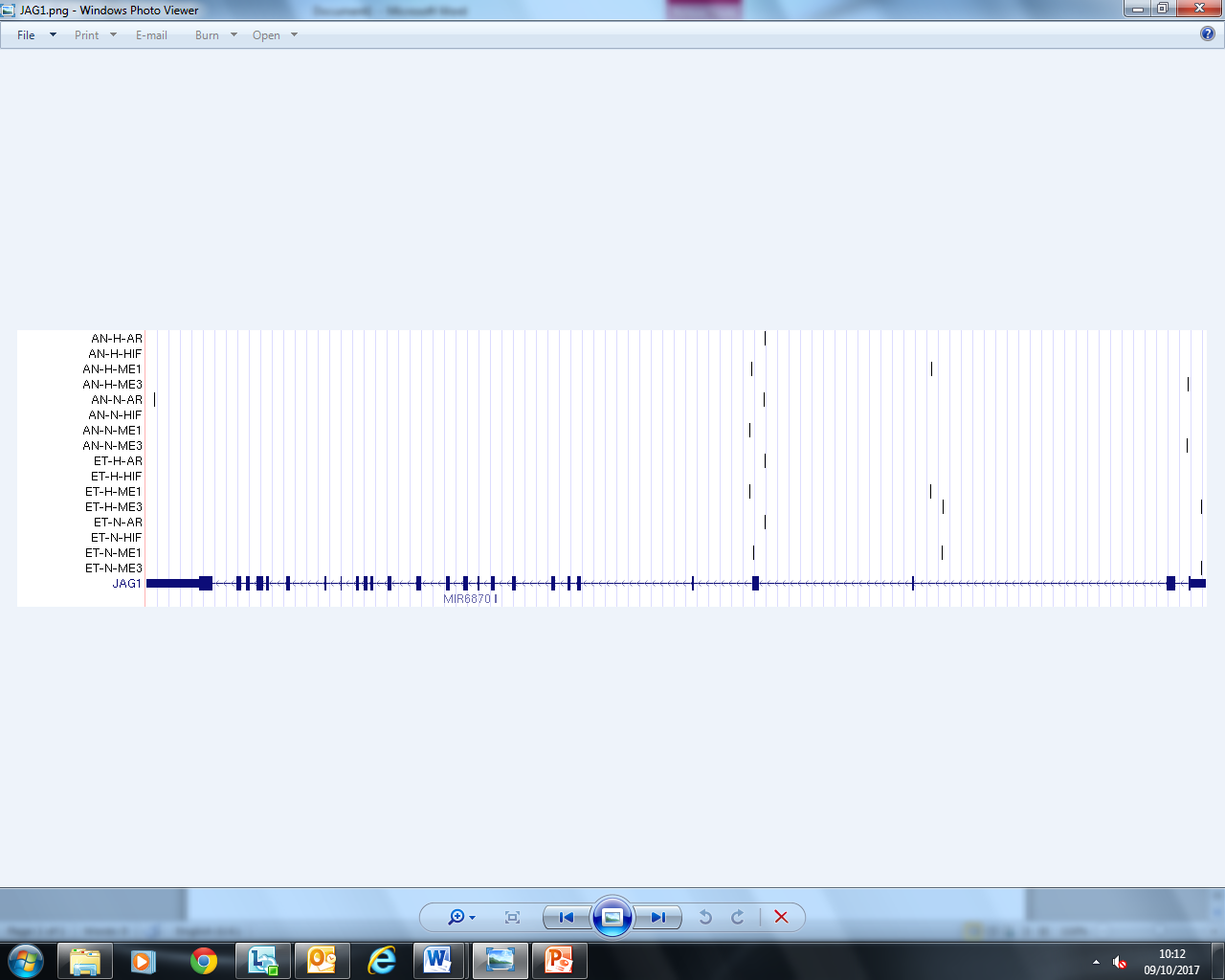

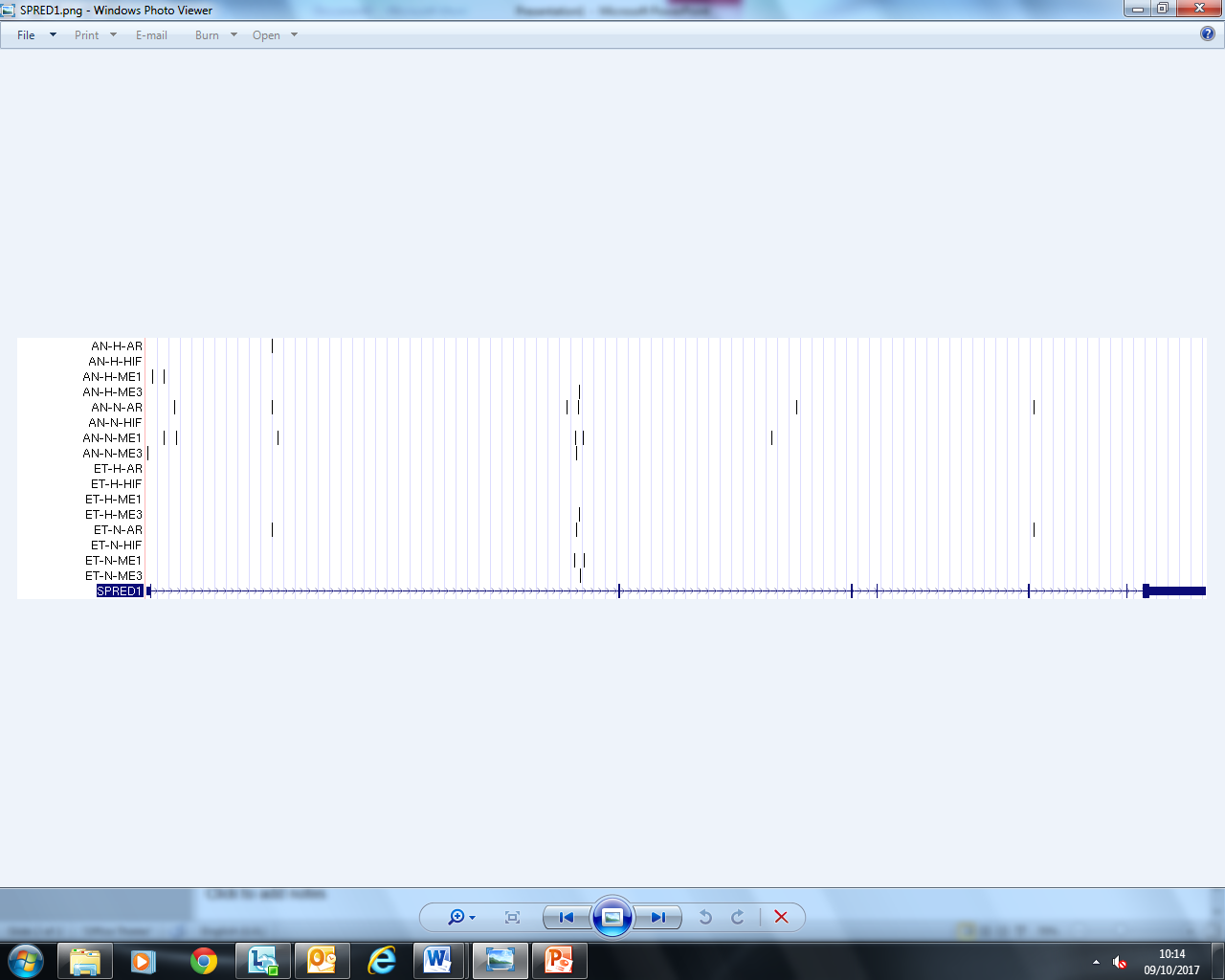

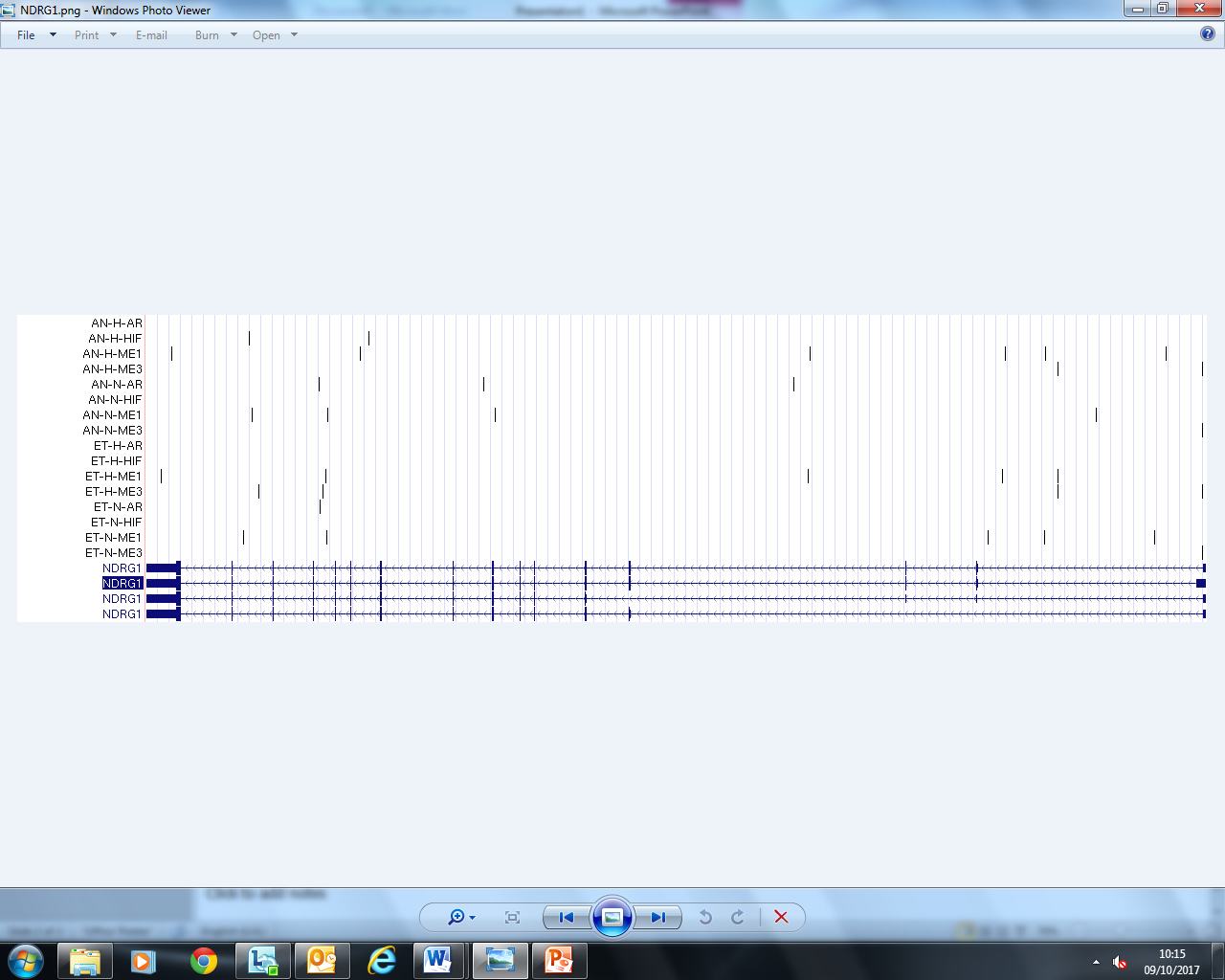

**Supplementary Figure 3.** HIF, AR, H3K4me1 and H3K4me3 binding sites within the introns (lines and arrows) and exons (boxes) of KCNN2, PPFIBP2, JAG1, SPRED1 and NDRG1. The ChIP-seq was performed using the LNCaP cell line following exposure to R1881 or ethanol under normoxia or hypoxia. The vertical lines represent the binding sites of the antibodies used to detect the annotated transcription factor or histone marker. AN; androgen R1881 treatment, ET; ethanol vehicle control, N; normoxia, H; hypoxia, ME1; H3K4me1 and ME3; H3K4me3.

**Supplementary Table I.** Publically available gene expression cohorts used in this study.

| Author | GEO Accession Number | Gene Expression Platform | Sample |
| --- | --- | --- | --- |
| TCGA [39] | N/A | Illumina RNA-seq | Surgical tissue |
| Taylor et al [16] | GSE21032 | Affimetrix Human Exon Arrays | Surgical tissue |
| Long et al [17] | GSE54460 | Illumina HiSeq | Surgical tissue |
| Sboner et al [18] | GSE16560 | Human 6k transcriptionally informative gene panel for DASL | Surgical tissue |
| Ross Adams et al [15] | GSE70770 | Illumina HT12 v4 BeadChip arrays | Surgical tissue |
| Massie et al [1] | GSE18684 | Illumina Human WG v2 BeadArray | Cell lines |

| GO Reference | Description | Number of Genes | Genes |
| --- | --- | --- | --- |
| GO:0045669 | Positive regulation of osteoblast differentiation | 8 | WNT7B, CEBPB, CEBPD, NPPC, JAG1, BMPR1B, BMPR1A, CYR61 |
| GO:0070059 | Intrinsic apoptotic signaling pathway in response to endoplasmic reticulum stress | 7 | TNFRSF10B, CEBPB, CHAC1, ERN1, TRIB3, PMAIP1, DDIT3 |

**Supplementary Table II.** GO and KEGG analysis of the 336 genes upregulated by androgen.

**Supplementary Table III.** GO and KEGG analysis of the 579 hypoxia upregulated genes.

| GO Reference | Description | Number of Genes | Genes |
| --- | --- | --- | --- |
| GO:0006351 | Transcription, DNA-templated | 87 | CREBRF, ZNF821, ZNF296, PPARG, CBX4, CBX2, RORA, MXI1, CBX8, ZNF16, GTF2IRD2B, GATA2, SAP30, CIPC, ZNF350, CGGBP1, KDM5B, SERTAD2, KMT5B, ELMSAN1, ZBTB21, ZFX, ARID5A, GTF2IRD2, NCOA7, IRF2BP2, TLE1, HES6, LPIN2, DAPK3, TOX2, DDIT3, ZFP37, EYA3, ASCL2, MAPK1, HES4, MNX1, TGIF1, PIAS2, VGLL4, ZNF586, CRTC3, ING2, CIART, FHL2, TRIB3, ZNF511, CXXC5, ZNF654, ZNF653, HINFP, CASZ1, BCL3, BCL6, BCL9L, KDM3A, HBP1, BHLHE40, ENO1, KLF6, ERF, CEBPB, CREBZF, KLF10, ZMYM5, TEAD2, HNRNPDL, MRGBP, POLR3C, BIRC2, TET1, FOXP1, UIMC1, ZNF165, RLF, ATXN1, PHF3, PHF1, NUPR1, ID1, IRF6, JMJD6, EBF2, KDM4B, PHF21A, ID3 |
| GO:0045944 | Positive regulation of transcription from RNA polymerase II promoter | 53 | NAMPT, ELF3, PPP2R5B, ELF4, ZNF821, PPARG, RORA, FOS, ACVR1B, GATA2, ZNF350, YAP1, TNIP1, CYR61, EGFR, ARHGEF2, NCOA7, DLL1, GPER1, LPIN2, DDIT3, JUN, NCK1, VEGFA, UBC, NFE2L1, PIAS2, MAPK7, CRTC3, SOX4, EGLN1, ATF2, SQSTM1, BCL3, KDM3A, BCL9L, DDX41, FOXD1, MAFF, CEBPB, CEBPG, ARID3A, TEAD2, TET1, FOXP1, RLF, ATF4, ATF3, CSRNP1, EBF2, IRF1, RBPJ, KLF4 |
| GO:0000122 | Negative regulation of transcription from RNA polymerase II | 43 | CREBRF, USP3, EFNA1, ZNF296, PPARG, CBX4, FHL2, TRIB3, CBX2, CXXC5, CBX8, ATF2, GATA2, SAP30, ZNF350, TSC22D3, CGGBP1, SQSTM1, YEATS2, HINFP, BCL6, BHLHE40, NFIL3, ERF, KLF10, KLF11, TLE1, HES6, FOXP1, DDIT3, FNIP1, ASCL2, ATF3, ID1, VEGFA, UBC, MNT, PHF21A, TGIF1, ID3, RBPJ, KLF4, VLDLR |
| GO:0045892 | Negative regulation of transcription, DNA-templated | 37 | ELF3, CIART, PPARG, CBX4, FHL2, TRIB3, TRIM11, CIPC, ZNF350, HINFP, YEATS2, BCL3, BCL6, BHLHE40, KDM5B, FOXD1, ENO1, CEBPB, ZBTB21, CREBZF, KLF10, ARID5A, KLF11, TLE1, UIMC1, FOXP1, DDIT3, ATXN1, CDKN1B, ID1, JUN, IRF1, ID3, RBPJ, RASD1, RBM15, KLF4 |
| GO:0043065 | Positive regulation of apoptotic processes | 28 | CDK19, LDHA, ING2, TNFRSF12A, SAV1, BNIP3, SOX4, SQSTM1, HMOX1, BCL6, FAM162A, ARHGEF2, KLF11, HRK, GPER1, DAPK3, JMY, ATF4, DUSP1, NUPR1, BNIP3L, UBC, SIAH1, ID3, GADD45B, GADD45A, IGFBP3, CAMK1D |
| GO:0008285 | Negative regulation of cell proliferation | 27 | ING2, SOX4, MXI1, PTEN, SPRY1, CDKN2C, CDKN2D, BCL6, NDRG1, ING1, TES, CGRRF1, KLF10, KLF11, DLL1, PIM2, GPER1, CDKN1B, BTG1, IRF6, JUN, MNT, IRF1, DNAJB2, RBPJ, IGFBP3, KLF4 |
| **GO:0001666** | **Response to hypoxia** | **20** | **LDHA, CLDN3, BNIP3, PDLIM1, EGLN1, BIRC2, DDIT4, ASCL2, PKM, VEGFB, LONP1, CDKN1B, PLOD1, CXCR4, ANG, PLOD2, HMOX1, VEGFA, HSD11B2, ALKBH5** |
| GO:0061621 | Canonical glycolysis | 15 | ALDOA, PFKL, PFKFB4, PFKFB3, ALDOC, HK2, PGAM1, PFKP, HK1, PKM, GPI, ENO2, PGK1, GAPDH, ENO1 |
| GO:0007050 | Cell cycle arrest | 15 | CGRRF1, TBRG4, RRAGD, DDIT3, JMY, CDKN1B, IRF6, CDKN2C, RASSF1, CDKN2D, ILK, IRF1, HBP1, PPP1R15A, GADD45A |
| GO:0006096 | Glycolytic process | 14 | ALDOA, LDHA, PFKL, ALDOC, HK2, PGAM1, HK1, GPI, PGAM4, PGM1, ENO2, PGK1, GAPDH, ENO1 |
| GO:0030308 | Negative regulation of cell growth | 13 | ACVR1B, CDKN1B, BTG1, CDKN2C, CDKN2D, PSRC1, PPARG, BCL6, DNAJB2, SESN2, ING1, ENO1, SERTAD2 |
| GO:0006094 | Gluconeogenesis | 12 | ALDOA, GPI, ATF4, ATF3, ALDOC, PGAM4, PGM1, ENO2, PGAM1, PGK1, GAPDH, ENO1 |
| **GO:0071456** | **Cellular response to hypoxia** | **12** | **ZFP36L1, VASN, STC2, HMOX1, VEGFA, BNIP3L, BNIP3, STC1, FAM162A, NDRG1, RORA, PTEN** |
| GO:0043433 | Negative regulation of sequence-specific DNA binding transcription factor activity | 10 | ID1, CEBPG, HMOX1, PIM1, PIAS2, EGLN1, BHLHE40, ID3, FLNA, DDIT3 |
| GO:0070373 | Negative regulation of ERK1 and ERK2 cascade | 9 | SPRY1, ATF3, DUSP1, PTPRR, GPER1, ERRFI1, PTEN, TNIP1, KLF4 |
| GO:0006950 | Response to stress | 9 | EGFR, MAPK1, CGRRF1, SQSTM1, DUSP10, HILPDA, GADD45B, ERRFI1, GADD45A |
| GO:0071276 | Cellular response to cadmium ion | 8 | AKR1C3, MT1A, HMOX1, MT1E, MT1H, MT1X, MT1G, MT1F |
| GO:0045926 | Negative regulation of growth | 6 | MT1A, MT1E, MT1H, MT1X, MT1G, MT1F |
| GO:0071294 | Cellular response to zinc ion | 6 | MT1A, MT1E, MT1H, MT1X, MT1G, MT1F |
| KEGG  Reference | Description | Number of Genes | Genes |
| Hsa04010 | MAPK signaling pathway | 25 | DUSP10, MAP4K2, MKNK2, FGF13, CACNB3, ATF2, FOS, MAX, MAPT, MAP3K1, HSPA6, PPP3CC, EGFR, PTPRR, FLNA, DDIT3, DUSP5, MAPK1, ATF4, DUSP1, JUN, HSPB1, MAPK7, GADD45B, GADD45A |
| Hsa01130 | Biosynthesis of antibiotics | 21 | BCKDHA, ALDOA, ODC1, LDHA, PFKL, ALDOC, PFKP, HK2, PGAM1, HK1, ACLY, AK4, PKM, GPI, CTH, PGAM4, PGM1, ENO2, PGK1, GAPDH, ENO1 |
| Hsa00010 | Glycolysis and gluconeogenesis | 17 | ALDOA, MINPP1, LDHA, PFKL, ALDOC, HK2, PGAM1, PFKP, HK1, PKM, GPI, PGAM4, PGM1, ENO2, PGK1, GAPDH, ENO1 |
| Hsa01200 | Carbon metabolism | 15 | ALDOA, PFKL, ALDOC, HK2, PGAM1, PFKP, HK1, PKM, GPI, PGAM4, ENO2, PGK1, GPT2, GAPDH, ENO1 |
| Hsa05230 | Central carbon metabolism in cancer | 13 | EGFR, PFKL, PFKP, HK2, PGAM1, HK1, SLC7A5, PTEN, PKM, SLC16A3, MAPK1, PGAM4, SLC2A1 |
| Hsa01230 | Biosynthesis of amino acids | 13 | ALDOA, PFKL, ALDOC, PFKP, PGAM1, PKM, CTH, PGAM4, ENO2, PGK1, GAPDH, GPT2, ENO1 |
| **Hsa04066** | **HIF-1 signaling pathway** | **13** | **EGFR, MAPK1, CDKN1B, PFKFB3, VEGFA, SLC2A1, MKNK2, ENO2, HK2, HK1, EGLN1, GAPDH, ENO1** |
| Hsa00051 | Fructose and mannose metabolism | 9 | ALDOA, MPI, PFKL, PFKFB4, PFKFB3, ALDOC, HK2, PFKP, HK1 |
| Hsa00500 | Starch and sucrose metabolism | 9 | GPI, GBE1, PYGL, PGM1, HK2, HK1, AMY1C, AMY1B, AMY1A |
| Hsa04978 | Mineral absorption | 9 | MT1A, SLC30A1, HMOX1, MT1E, SLC31A1, MT1H, MT1X, MT1G, MT1F |
| Hsa00030 | Pentose phosphate pathway | 6 | ALDOA, GPI, PFKL, ALDOC, PGM1, PFKP |
